# Production of mouse ultrasonic vocalizations and distress calls is associated with different patterns of Fos expression in the nucleus retroambiguus

**DOI:** 10.64898/2026.03.18.712517

**Authors:** Patryk Ziobro, Da-Jiang Zheng, Aditya Rawal, Zizhan Zhou, Asmita Mittal, Katherine Tschida

## Abstract

Animals produce different vocalization types, which differ in their acoustic features and are produced in different behavioral contexts. How vocalization-related brain circuits are organized to enable the production of different vocalization types remains poorly understood. The nucleus retroambiguus is a hindbrain premotor region that regulates the production of both ultrasonic vocalizations (USVs) and distress calls (squeaks) in adult mice, but whether distinct or overlapping populations of RAm neurons are recruited during the production of these two vocalization types is unknown. In the current study, we used Fos immunohistochemistry to compare the counts and spatial distributions of Fos-positive RAm neurons in males and females that produced USVs and females that produced courtship squeaks. We also combined in vivo activity-dependent (TRAP2) labeling with Fos immunohistochemistry to directly compare Fos expression associated with the production of USVs and courtship squeaks in the same females. Our findings suggest that RAm contains three vocalization-related populations of neurons: squeak-related neurons, USV-related neurons, and shared neurons that are recruited during both vocalization types. These findings refine current models of the premotor control of vocalization and set the stage for future work to explore anatomical and functional heterogeneity within RAm.

## Introduction

To communicate successfully, animals produce different vocalization types, which differ in their acoustic features and are produced in different behavioral contexts (e.g., contact calls, alarm calls, mating calls, etc.). In mammals, vocalization requires an increase in expiratory airway pressure coordinated with the adduction of laryngeal muscles, and the resulting sound can be further influenced by vocal tract filtering. Thus, despite their differences in acoustic features, the production of different vocalization types requires the recruitment of overlapping sets of laryngeal, respiratory, and orofacial muscles. How vocalization-related brain circuits are organized to enable the production of different vocalization types remains poorly understood.

The nucleus retroambiguus (RAm) is a hindbrain region that plays a well-established role in vocal production. RAm is situated in the intermediate reticular formation caudal to the nucleus ambiguus and extends caudally to the brain-spinal cord transition (Jürgens, 2002, 2009). RAm sends axonal projections to the nucleus ambiguus, which houses laryngeal motor neurons, as well as to the thoracic and lumbar spinal cord, which houses intercostal and abdominal motor neurons (Miller et al., 1985; Merrill and Lipski, 1987; Holstege, 1989; Katada et al., 1996). RAm also receives a robust descending projection from the lateral periaqueductal gray (PAG), a midbrain region that gates the production of innate vocalizations in mammals (Jürgens, 1994, 2002, 2009). Aside from its anatomical positioning, substantial functional evidence implicates RAm in the regulation of vocal production. Artificial activation of RAm-projecting PAG neurons elicits the production of vocalizations in mice (Tschida et al., 2019; Park et al., 2024), and work in decerebrate cats found that transections of the medulla at the level of RAm, as well as excitotoxic lesions of the reticular formation that include RAm, block PAG-elicited vocalizations (Zhang et al., 1995; Shiba et al., 1997). Conversely, direct pharmacological stimulation of RAm elicits the production of vocalizations in decerebrate cats, albeit with special-atypical acoustic features (Zhang et al., 1992, 1995; Subramanian and Holstege, 2009). Together, these past studies support an important role for RAm in regulating vocal production but leave open the question of how RAm participates in the production of different vocalization types.

Two recent studies in mice have begun to shed light on this question. Mice produce two main vocalization types: (1) ultrasonic vocalizations (USVs), which are produced by adults during social interactions (Nyby, 1983; Holy and Guo, 2005; Moles et al., 2007; Warren et al., 2020; Zhao et al., 2021, 2025) and by mouse pups when isolated from the nest (Okon, 1970; Noirot, 1972; Ehret, 2005; Pranic et al., 2022), and (2) distress calls (squeaks), which are produced in aversive behavioral contexts (Grimsley et al., 2013, 2016; Finton et al., 2017; Ruat et al., 2022; Ziobro et al., 2024). The first study used an activity-dependent labeling strategy to express viral transgenes in RAm neurons that upregulated expression of the immediate early gene Fos following the production of courtship USVs in male mice (Park et al., 2024). Expression of tetanus toxin light chain in these “RAm-voc” neurons subsequently blocked the production of both male courtship USVs and squeaks elicited by tail pinch or footshock, suggesting that at least some RAm neurons regulate the production of both vocalization types. However, optogenetic activation of RAm-voc neurons elicited only USV production, consistent with the idea that the activity of additional neurons (either within RAm or elsewhere) is needed to drive the production of squeaks. A second study found that a large proportion of neurotensin (*Nts*)-expressing RAm neurons are Fos-positive after the production of isolation USVs in pups, as well as after the production of courtship USVs in adult males (Veerakumar et al., 2023). Caspase-mediated ablation of RAm *Nts* neurons blocked the production of male courtship USVs. Notably, optogenetic activation of RAm *Nts* neurons at low frequencies elicited the production of squeak-like (< 20 kHz) vocalizations, while optogenetic activation at higher frequencies elicited the production of vocalizations in the ultrasonic range. Together, these studies demonstrate that RAm regulates the production of both USVs and squeaks, but the degree to which overlapping or distinct sets of RAm neurons are recruited during the two vocalization types remains unclear. Addressing this question requires a direct comparison of RAm neuronal activation associated with the production of USVs and squeaks.

In the current study, we used immunohistochemistry in a between-subjects design to compare the counts and spatial distributions of Fos-positive RAm neurons following the production of USVs in adult males and females vs. following the production of squeaks during courtship interactions (females only). We also directly compared RAm Fos expression associated with the production of USVs and courtship squeaks in females by combining in vivo activity-dependent labeling (TRAP2) with Fos immunohistochemistry in a within-subjects design. Our findings refine existing models of premotor control of different vocalization types and highlight the need for work to characterize anatomical and functional heterogeneity in vocalization-related RAm neurons.

## Methods

### Ethics statement

All experiments and procedures were conducted according to protocols approved by the Cornell University Institutional Animal Care and Use Committee (protocol #2020-001).

### Subject details

Male and female TRAP2;Ai14 mice and C57 mice (Jackson laboratories, 000664) were housed with their siblings and both parents until weaning at postnatal day 21. TRAP2;Ai14 mice were generated by crossing TRAP2 mice (Jackson Laboratories, 030323) with Ai14 mice (Jackson Laboratories, 007914). All mice were kept on a 12:12 reversed light/dark cycle, were housed in ventilated micro-isolator cages in a room with regulated temperature and humidity, and were provided with unrestricted access to food and water. A running wheel (Innovive) was present in all home cages from the time of weaning and was subsequently removed immediately prior to any behavioral trials that were conducted in the subject mouse’s home cage.

### RAm Fos study design

Five groups of C57 mice were subjected to different treatments prior to collecting brains and quantifying Fos expression in RAm: (1) males that produced USVs during courtship interactions with females; (2) females that produced USVs during same-sex interactions with females; (3) females that produced squeaks during courtship interactions with males; (4) males that produced squeaks during a mild footshock paradigm (see below for details of footshock delivery); (5) and females that produced squeaks during a mild footshock paradigm. Subject mice in the male USV, female USV, and female courtship squeak groups were single-housed for 3-days prior to the behavioral trial. The majority of social interaction trials were 30 minutes in duration, and a subset were shorter (1-10 minutes in duration) to elicit lower vocalization rates from subject mice. For subjects in the male USV and female USV groups, the subject mouse’s home cage was placed in a sound-attenuating recording chamber (Med Associates) equipped with an ultrasonic microphone (Avisoft), an infrared light source (Tendelux), and a webcam (Logitech, with the infrared filter removed to enable video recording under infrared lighting conditions). A novel, group-housed female was then placed in the home cage of the subject mouse, and vocal and non-vocal behaviors were recorded. For subjects in the female courtship squeak group, the female was placed into the home cage of a novel male inside a sound-attenuating recording chamber, with the bedding removed to facilitate the detection of courtship squeaks. At the end of behavior trials, subject mice remained or were returned to their home cage and were sacrificed 120 minutes from the start of the behavior trial.

The sample sizes in the five groups were as follows: male USV (n = 12), female USV (n = 16), female courtship squeak (n = 20), male footshock (n = 4), and female footshock (n = 3).

### RAm overlap study design

One group of male TRAP2;Ai14 mice (male USV-USV group) and four groups of female TRAP2;Ai14 mice (female USV-USV, female squeak-squeak, and female squeak-USV, and female USV-squeak groups) were included in this study. Subject mice were single-housed (day 0), and three days later (day 3) were subjected to social interactions to elicit high rates of either USVs or courtship squeaks (females only). Males and females in the USV-USV groups were given 30-minute-long interactions with novel females to elicit high rates of USV production, and females in the squeak-squeak and squeak-USV groups were given 30-minute-long courtship interactions with males to elicit the production of high rates of courtship squeaks, as described above. Following these social interactions, subject mice received I.P. injections of 4-OHT (50 mg/kg) to enable the expression of tdTomato in recently active neurons, including in RAm neurons that express Fos in association with the production of USVs/squeaks. Ten days later (day 13), subject mice were given either a 30-minute social interaction to elicit USVs (male USV-USV, female USV-USV, and female squeak-USV groups) or a 30-minute courtship interaction to elicit squeaks (female squeak-squeak and female USV-squeak groups). At the end of this second behavior trial, subject mice remained or were returned to their home cage and were sacrificed 120 minutes from the start of the behavior trial. The sample sizes in the five groups were as follows: male USV-USV (n = 5), female USV-USV (n = 6), female squeak-squeak (n = 6), female squeak-USV (n = 5), and female USV-squeak (n = 6).

### Drug preparation

4-hydroxytamoxifen (4-OHT, HelloBio, HB6040) was dissolved at 20 mg/mL in ethanol by shaking at 37°C and was then aliquoted (75 uL) and stored at -20°C. Before use, 4-OHT was redissolved in ethanol by shaking at 37°C and filtered corn oil was added (Sigma, C8267, 150 uL). Ethanol was then evaporated by vacuum under centrifugation to give a final concentration of 10 mg/mL, and 4-OHT solution was used on the same day it was prepared, delivered at a final dosage of 50 mg/kg.

### Immunohistochemistry

Two hours following the start of behavior trial, subject mice were deeply anesthetized using isoflurane and then transcardially perfused with phosphate-buffered saline (PBS, pH 7.4), followed by 4% paraformaldehyde (PFA, in 0.1 M PBS, pH 7.4). Brains were subsequently dissected and post-fixed in 4% PFA for 24 hours at 4°C, followed by immersion in 30% sucrose solution in PBS for 48 hours at 4°C. Afterward, brains were embedded in frozen section embedding medium (Surgipath, VWR), flash frozen in a dry ice-ethanol (100%) bath, and then stored at ^−^80°C until sectioning. Coronal sections containing RAm were cut on a cryostat to a thickness of 80 μm, washed in PBS (3 x 5 mins at RT), permeabilized for 2-3 hours in PBS containing 1% Triton X-100 (PBST), and then blocked in 0.3% PBST containing 10% Blocking One (Nacalai USA) for 1 hour at RT on a shaker. Sections were then incubated for 24 hours at 4°C with primary antibody in blocking solution (1:1000 rabbit anti-Fos, Cell Signaling Technologies, 2250S), washed 3 × 10 minutes in 0.3% PBST, then incubated for 24 hours at 4°C with secondary antibody in blocking solution (1:1000, Alexa Fluor 488 goat-anti-rabbit, Invitrogen, plus 1:500 NeuroTrace, Invitrogen). Finally, sections were washed for 2 × 10 minutes in 0.3% PBST, followed by washing for 2 × 10 minutes in PBS. After mounting on slides, sections were dried and coverslipped with Fluromount G (Southern Biotech). Slides were imaged with a 10x objective on a Zeiss 900 laser scanning confocal microscope. To generate the images in Fig. 1A, immunostaining for Fos was combined with immunostaining for choline acetyltransferase (ChAT) (these cases were not used for Fos quantification). In these cases, sections were incubated with primary antibodies in blocking solution (1:1000 rabbit anti-Fos and 1:1000 goat anti-ChAT, Millipore, AB144P) and later with secondary antibodies (1:1000 Alexa Fluor 488 donkey anti-rabbit, Invitrogen, and 1:1000 Alexa 594 donkey anti-goat, Invitrogen) but were otherwise processed as described above.

**Figure 1.**
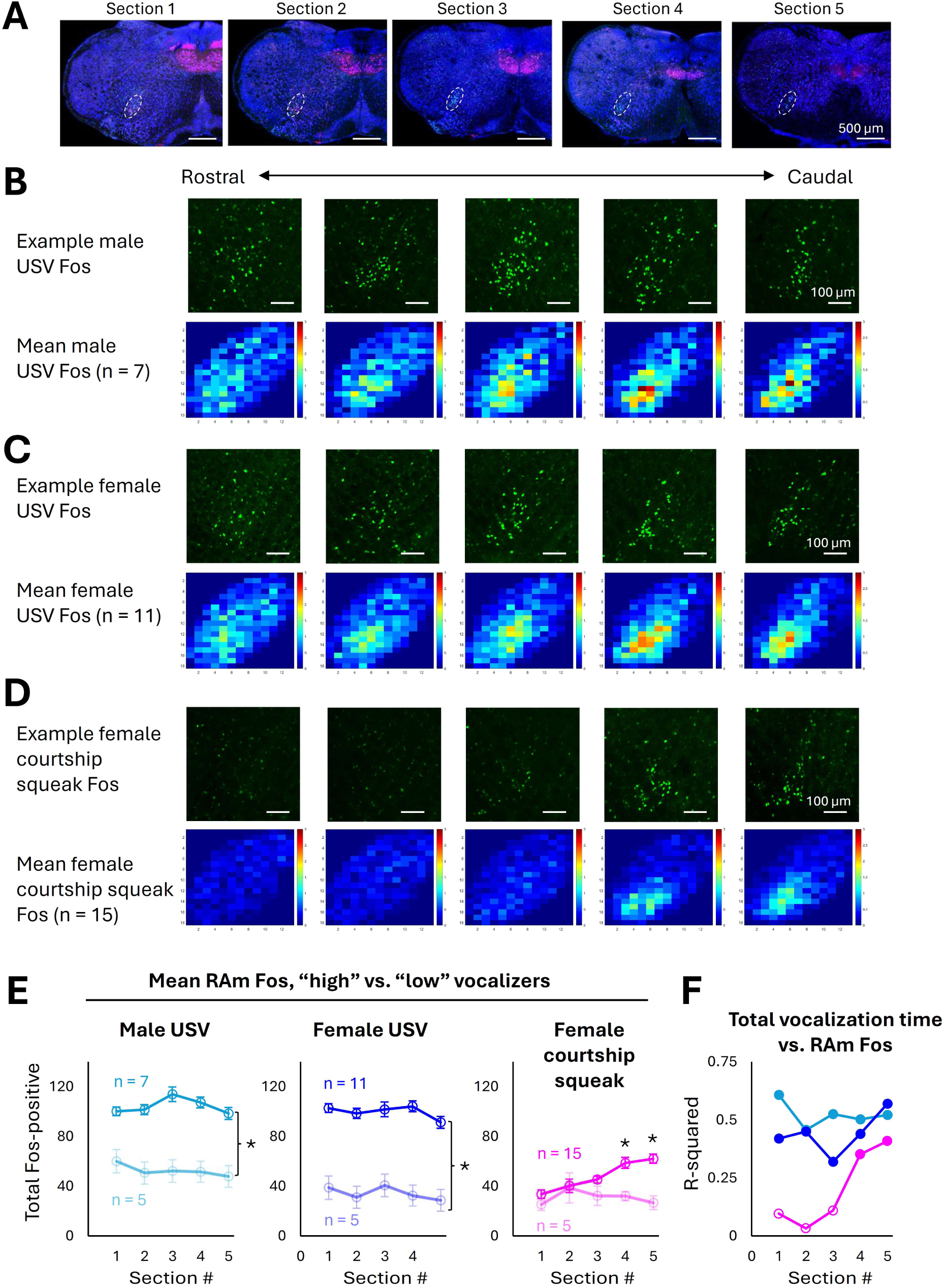
Production of USVs vs. squeaks drives different patterns of Fos expression in RAm. (A) Representative coronal sections show the extent of RAm Fos expression associated with male USV production. White dotted ovals show ROI placement for Fos scoring. Blue, Neurotrace; green, Fos; red, choline acetyltransferase. (B) Representative confocal images (top row) show RAm Fos expression in a male that produced USVs during a 30-minute courtship interaction with a female. Bottom row shows mean heatmap representations for the high male USV group. (C) Same as (B), for RAm Fos expression in females that produced USVs during 30-minute same-sex interactions with females. (D) Same as (B), for RAm Fos expression in females that produced courtship squeaks during 30-minute interactions with males. (E) Mean counts of Fos-positive RAm neurons are shown by section (1-5) in high vs. low USV males (left), high vs. low USV females (middle), and high vs. low courtship squeak females (right). (F) Plot shows the R-squared values for the correlation between total Fos-positive RAm neurons and total vocalization time for each group and for each RAm section. Filled points indicate statistically significant relationships, and open points indicate non-significant relationships.

### Confocal image analyses

RAm is located caudal to the nucleus ambiguus in the intermediate reticular formation. In this study, we defined the boundaries of RAm according to the extent of USV-associated Fos expression observed in male mice. For confocal images that included USV-associated RAm Fos and/or tdTomato expression, five coronal sections were selected that included RAm at the following approximate A/P positions: -7.2 mm, -7.4 mm, -7.6 mm, -7.8 mm, -8.0 mm caudal to bregma. In each plane of section, oval-shaped regions of interest (ROIs, 200 pixels wide by 400 pixels tall) were generated in ImageJ, rotated 30 degrees (plus 30 degrees for left RAm, minus 30 degrees for right RAm), and then were placed over the bands of Fos-positive and/or tdTomato-positive neurons in the intermediate reticular formation that correspond to RAm. For confocal images from mice that did not include USV-associated RAm Fos or tdTomato, anatomical landmarks (inferior olive, lateral reticular nucleus, spinal trigeminal nucleus) were used to match the placement of RAm ROIs to the placement from images that included USV-associated RAm Fos/tdTomato. For mice in the RAm Fos study, Fos-positive neurons located within these RAm ROIs were manually annotated in each of the five sections by trained observers using ImageJ. For mice in the RAm overlap study, tdTomato-positive neurons located within these RAm ROIs were manually annotated, and overlap was calculated as the proportion of tdTomato-positive RAm neurons that also expressed Fos. Heatmaps of RAm Fos expression and double-labeled RAm neurons (Fos-positive and tdTomato-positive) were constructed as follows. The locations of annotated neurons in each coronal section were first standardized with respect to the position of their RAm ROI (i.e., neuron coordinates were calculated in pixels relative to the centroid of their RAm ROI, with right RAm neuron coordinates flipped horizontally in order to be pooled with the neuron coordinates from left RAm for that same plane of section). To create heatmaps for each plane of section for a single mouse, a bounding rectangle (266 pixels wide by 362 pixels tall) that encompasses the 200 x 400 pixel RAm ROI was created and organized into 13 horizontal bins and 18 vertical bins (each bin approximately 20 x 20 pixels). For a given plane of section, the standardized location of each annotated RAm neuron was then placed in its corresponding heatmap bin, and the total number of annotated RAm neurons in each heatmap bin was calculated. This procedure was repeated to generate a heatmap for each plane of section (sections 1-5) for each mouse. Mean heatmaps for sections 1-5 were then created for experimental groups by averaging the heatmaps generated for individual mice in that group for each plane of section. Please note that heatmap representations only reflect Fos quantification within the RAm ROIs, and tdTomato-positive and/or Fos-positive neurons outside the RAm ROIs were not counted.

### USV recording and detection

USVs were recorded with an ultrasonic microphone (Avisoft, CMPA/CM16), amplified (Presonus TubePreV2), and digitized at 250 kHz (Spike 7, CED). USVs were detected with custom Matlab codes (Tschida et al., 2019) using the following parameters: mean frequency > 45 kHz; spectral purity > 0.3; spectral discontinuity < 1.0; minimum USV duration = 5 ms; minimum inter-syllable interval = 30 ms).

### Footshock delivery

Subject mice were placed in a footshock chamber (Med Associates) inside a sound attenuating chamber (Med Associates) equipped with an ultrasonic microphone (Avisoft) and a webcam (Logitech). A mild (0.5 mA, 1.5s-long) shock was delivered once every 30 seconds, for a total of 10 footshocks (trial duration = 5 minutes).

### Squeak detection

In footshock and courtship trials (females only), audio recordings were generated as described above for USV recordings, and a custom Matlab code was used that allowed trained users to manually annotate the onsets and offsets of individuals squeaks from spectrograms created from audio recordings. The total number of squeaks and the total squeak time were calculated from these annotations.

### Statistics

The Shapiro-Wilk test was used to evaluate the normality of data distributions. Parametric, two-sided statistical comparisons were used in all analyses of normally distributed data, and non-parametric, two-sided comparisons were used in analyses that included non-normally distributed data (alpha = 0.05). No statistical methods were used to pre-determine sample size. Errors represent standard error of the mean unless otherwise noted.

### Data availability

Source data generated in this study, as well as custom Matlab codes, are available upon request.

## Results

### Production of ultrasonic vocalizations (USVs) vs. courtship squeaks drives different patterns of Fos expression in the nucleus retroambiguus (RAm)

As a first step toward comparing Fos expression in RAm associated with the production of USVs vs. squeaks, we characterized the rostral-to-caudal extent of RAm Fos expression associated with USV production in male mice. Although past studies have reported that RAm neurons upregulate Fos expression in males that recently produced high rates of courtship USVs (Tschida et al., 2019; Concha-Miranda et al., 2022; Veerakumar et al., 2023; Park et al., 2024), the spatial extent of USV-associated RAm Fos expression has not been described. To this end, male subjects were single-housed for 3 days to promote high rates of USV production and then were given 30-minute social interactions with novel, group-housed females. Males were perfused 120 minutes later, and hindbrain sections were collected and stained for Fos expression, starting at the level of the nucleus ambiguus and extending caudally to the hindbrain-spinal cord transition. We found that male USV production drove robust Fos expression in neurons within the intermediate reticular formation, beginning around -7.2 mm caudal to bregma and extending caudally approximately 1 mm (n = 7 high USV males; Fig. 1A-B, E; see Methods for details of heatmap representations). To better evaluate the relationship between RAm Fos expression and male USV production, we also added behavioral data and measured RAm Fos expression from n = 5 ‘low USV’ males that were given brief (1-10 minute) social interactions in order to elicit lower USV rates. High vs. low USV males differed significantly in the total numbers of Fos-positive neurons across all 5 RAm sections (Fig. 1E; F(1,10) = 17.03, p = 0.002 for main effect of group, p > 0.05 for main effect of section and for interaction effect; two-way ANOVA with repeated measures on one factor). When we pooled high and low male USV cases to calculate the correlation between total vocalization (USV) time and total Fos-positive neurons for each RAm section (1-5), we found that total USV time was significantly correlated with RAm Fos expression across all 5 RAm sections in the male USV group (Fig. 1F, Fig. S1; p < 0.05 for all male USV comparisons; linear regressions). We subsequently defined the boundaries of RAm according to this male USV-associated pattern of Fos expression (see Methods).

We next asked whether USV production in female mice was associated with a similar pattern of RAm Fos expression. To elicit robust USV production from females, 3-days-single-housed female subjects were given 30-minute social interactions with novel females. Although we did not employ microphone array methods to assign individual USVs to the subject female vs. the novel female (Heckman et al., 2017; Warren et al., 2020; Matsumoto et al., 2022), past work found that short-term social isolation promotes multiple aspects of female social behavior during same-sex interactions (Zhao et al., 2021, 2025), and the observed high USV rates suggest that single-housed subject females vocalize robustly during these interactions (mean total female-female USVs = 2567 ± 292). Correspondingly, female subjects exhibited robust RAm Fos expression with the same localization seen in males (n = 11 high USV females; Fig. 1C). To evaluate the relationship between RAm Fos expression and female USV production, we added behavioral data and measured RAm Fos expression from n = 5 ‘low USV’ females that were given brief social interactions in order to elicit lower USV rates. Similar to the male USV group, high vs. low USV females differed significantly in the total numbers of Fos-positive neurons across all 5 RAm sections (Fig. 1E; F(1,14) = 85.36, p < 0.001 for main effect of group, p > 0.05 for main effect of section and for interaction effect; two-way ANOVA with repeated measured on one factor). Also like the male USV group, when we considered the correlation between total vocalization (USV) time and total Fos-positive neurons for each RAm section (1-5), we found that total USV time was significantly correlated with RAm Fos expression across all 5 RAm sections in the female USV group (Fig. 1F, Fig. S1; p < 0.05 for all female USV comparisons; linear regressions). In summary, USV production is associated with robust and similar patterns of RAm Fos expression in males and females.

Having characterized the spatial extent of RAm Fos expression associated with USV production, we next characterized RAm Fos expression associated with the production of squeaks. We initially considered RAm Fos expression in a small number of male and female mice that were subjected to a mild footshock paradigm to elicit squeaks (Fig. S2; see Methods; n = 3 female footshock, n = 4 male footshock). However, these mice produced only modest rates of squeaks (20-60 total squeaks), and the total number of Fos-positive RAm neurons was low relative to females and males that produced high rates of USVs (Fig. S2; mean total RAm Fos-positive neurons: male footshock = 93 ± 21, female footshock = 193 ± 29, female USV = 499 ± 15, male USV = 545 ± 36). In male mice, we are unaware of an alternative behavioral paradigm that would enable us to elicit substantially higher squeak rates without also risking injury to our subject animals (e.g., interactions with an aggressive male). As an alternative paradigm to elicit higher rates of squeaks in females, we measured female squeak rates during courtship interactions with males (i.e., courtship squeaks). We found that females produced robust rates of squeaks during 30-minute courtship interactions with sexually experienced males (Fig. S2C; mean total courtship squeaks = 233 ± 31; t(20) = 3.85, p < 0.001 for difference in total squeaks between courtship and footshock groups; t-test). When we examined RAm Fos expression in female mice that produced high rates of courtship squeaks, we found robust Fos expression, but in a pattern that differed in two ways from RAm Fos expression associated with USV production. First, courtship squeak production is associated with overall weaker RAm Fos expression than either male or female USV production (Fig. 1D; F(2,30) = 71.31, p < 0.001 for main effect of group, p < 0.0001 for male USV vs. female courtship squeak, p < 0.0001 for female USV vs. female courtship squeak; one-way ANOVA plus post-hoc Tukey’s HSD).

Second, we also observed a difference in the localization of squeak-associated RAm Fos expression. Whereas USV production was associated with robust Fos expression across all 5 representative planes of section, squeak production was associated with robust Fos expression in only the caudal-most RAm sections (sections 4-5; Fig. 1D). To evaluate the relationship between this pattern of RAm Fos expression and female courtship squeak production, we added behavioral data and measured RAm Fos expression from n = 5 ‘low courtship squeak’ females that were given brief social interactions with males in order to elicit lower squeak rates. Notably, the mean total counts of Fos-positive RAm neurons tended to be higher in RAm sections 4 and 5 but differed significantly only in the caudal-most RAm between high courtship squeak and low courtship squeak females (Fig. 1E; F(2,32) = 5.0, p = 0.008 for interaction between group and section, p = 0.004 for section 5 high squeak vs. low squeak; two-way ANOVA with repeated measures on one factor plus post-hoc Tukey’s HSD for across-group, within-section comparisons). Similarly, when we considered the correlation between total vocalization (squeak) time and RAm Fos expression for each RAm section, we found that total squeak time was only significantly correlated with RAm Fos expression in sections 4-5 (Fig. 1F, Fig. S1; p < 0.05 for section 4 and section 5 comparisons only; linear regressions). In summary, these data show that production of female/male USVs and female courtship squeaks are associated with different patterns of Fos expression in RAm.

One possibility is that the difference in counts and spatial distributions of Fos-positive RAm neurons associated with USVs vs. squeaks is driven by the production of different vocalization types per se. However, another possibility is that the difference in RAm Fos expression is driven by the difference in total vocalization time between the USV and courtship squeak groups.

Although we were able to elicit robust squeak rates using the female courtship squeak paradigm, females nonetheless spent on average less total time squeaking than males spent producing USVs (Fig. 2A; z = 3.67, p < 0.001, Mann Whitney U test; female USV group not included in comparison because the number of USVs produced by subject females vs. novel females was not determined). To test whether a difference in vocalization time could account for the observed difference in RAm Fos expression patterns, we conducted two analyses. First, we selected a subset (n = 4) of male USV cases that were well matched in their total vocalization time to a subset (n = 4) of female courtship squeak cases (Fig. 2A). A comparison of these cases showed that male USV production was still associated with greater number of Fos-positive RAm neurons than female courtship squeak production (Fig. 2B; F(1,6) = 13.28, p = 0.01 for main effect of group, p > 0.05 for main effect of section and for interaction effect; two-way ANOVA with repeated measures on one factor). Moreover, heatmap representations of the mean RAm Fos expression for the two groups revealed patterns of Fos expression that resemble Fos expression in the full male USV and full female courtship squeak groups (compare Fig. 2C vs. Fig. 1C-D).

**Figure 2.**
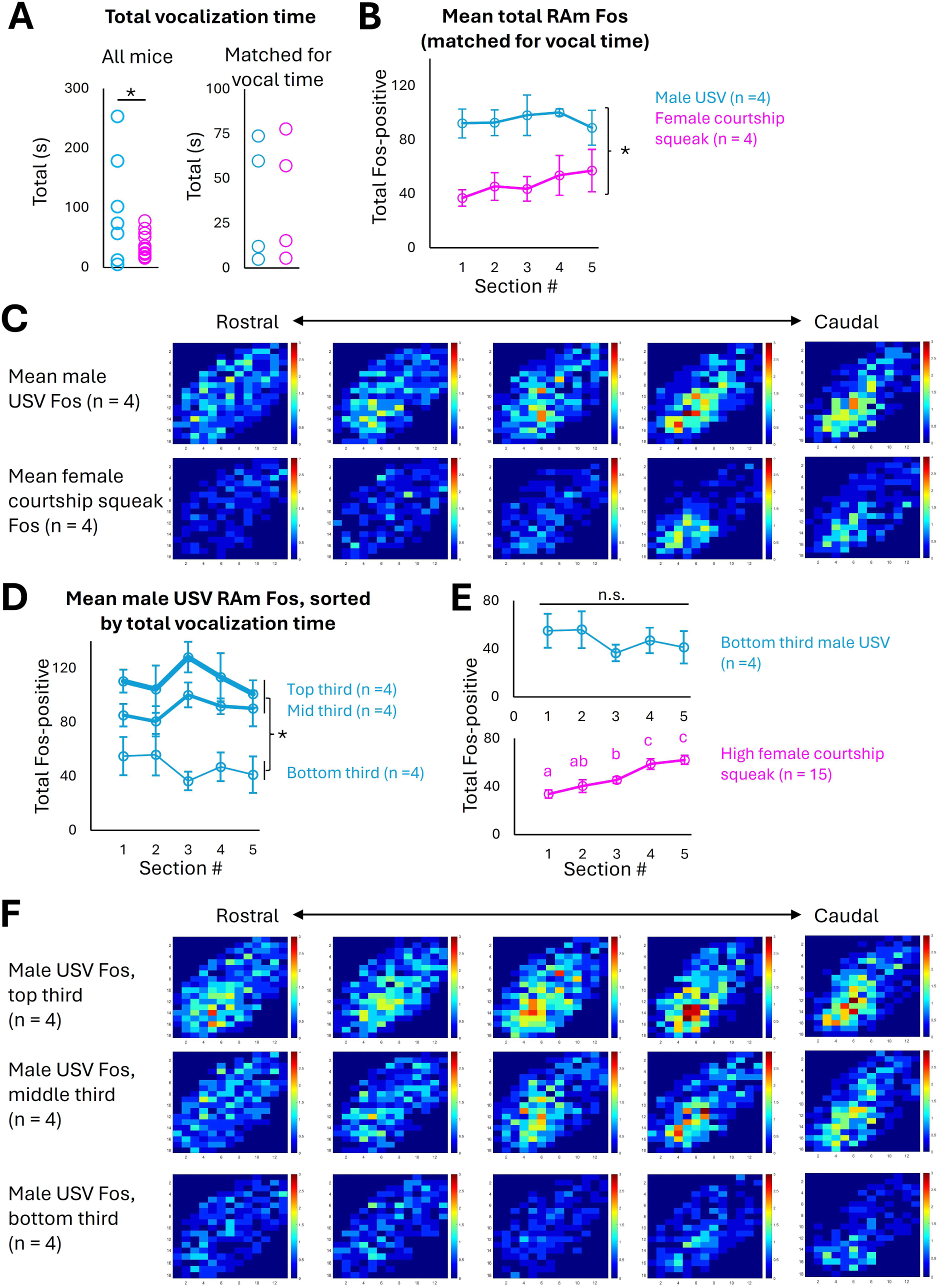
Differences in RAm Fos expression following the production of USVs vs. courtship squeaks are not accounted for by differences in total vocalization time. (A) (Left) Total vocalization time is plotted for all high male USV cases (n = 7, blue) and high female courtship squeak cases (n = 15, magenta). (Right) Total vocalization time is plotted for n = 4 high male USVs cases and n = 4 high female courtship squeak cases that are matched for vocalization time. (B) Mean counts of Fos-positive RAm neurons are plotted by section for the subset of male USV and courtship squeak cases matched for total vocalization time. (C) Heatmaps show mean RAm Fos expression by section for the subset of male USV and courtship squeak cases matched for total vocalization time. (D) Mean counts of Fos-positive RAm neurons are plotted by section for male USV cases split into groups according to total vocalization time. (E) Mean counts of Fos-positive RAm neurons are plotted by section for the bottom-third male USV cases (n = 4, top) and for the high female courtship squeaks cases (n = 15, bottom; replotted from Fig. 1E). (F) Heatmaps show mean RAm Fos expression by section for the top-third, middle-third, and bottom-third of male USV cases, grouped by total vocalization time.

Second, we split the male USV cases into three groups according to their total vocalization time (Fig. 2D-F; n = 4 males in the top-third, n = 4 males in the middle-third, and n = 4 males in the bottom-third USV groups). Total RAm Fos expression was higher in the top-third and middle-third male USV groups than in the bottom-third male USV group (Fig. 2D; F(2,9) = 10.17, p = 0.005 for main effect of group, p > 0.05 for main effect of section and for interaction effect; p = 0.004 for top vs. bottom, p = 0.04 for mid vs. bottom; two-way ANOVA with repeated measures on one factor plus post-hoc Tukey’s HSD). Even in the bottom-third male USV group, however, we found that the distribution of Fos-positive RAm neurons across sections 1-5 still looked “USV-like” rather than “squeak-like”. That is to say, in the high female courtship squeak group, RAm Fos expression is significantly higher in the caudal sections of RAm than in the rostral sections of RAm (Fig. 2E; magenta points show high female courtship squeak data replotted from Fig. 1E; F(2.16, 30.23) = 14.40, p < 0.001 for main effect of section; one-way ANOVA with repeated measures plus post-hoc Tukey’s HSD; different lower-case letters indicate significant differences between sections). In contrast, in the bottom-third male USV group, RAm Fos expression is not significantly different between the rostral and caudal sections of RAm (Fig. 2E; F(4,12) = 1.59, p > 0.05 for main effect of section; one-way ANOVA with repeated measures). Heatmap representations of mean RAm Fos expression also show that although RAm Fos expression is weakest in the bottom-third male USV group, Fos expression remains distributed across both rostral and caudal RAm sections (Fig. 2F). In summary, these analyses indicate that differences in total vocalization time cannot account for the different patterns of RAm Fos expression associated with the production of USVs and courtship squeaks. Rather, these differences support the conclusion that at least somewhat distinct sets of RAm neurons upregulate Fos following the production of USVs vs. squeaks.

### Only a subset of RAm neurons express Fos following the production of both USVs and squeaks

The Fos datasets indicate that squeak production is associated with Fos expression biased toward caudal RAm, while USV production is associated with broader Fos expression throughout rostral and caudal RAm. To what degree are RAm neurons recruited during both vocalization types? On the one hand, RAm neurons that are activated during squeak production could represent a nested subset of the RAm neurons that are activated during USV production, similar to the organization recently described for ventral nerve cord circuits that regulate the production of pulse and sine courtship song in *Drosophila* males (Lillvis et al., 2024; Shiozaki et al., 2024). Alternatively, RAm might contain USV-related neurons, squeak-related neurons, and shared neurons that are recruited during both vocalization types. To directly compare Fos expression associated with the production of USVs vs. squeaks in the same animals, we combined in vivo activity-dependent labeling (TRAP2) with Fos immunohistochemistry in a within-subjects design (Fig. 3A). Briefly, each TRAP2;Ai14 subject was exposed to a behavioral context that would elicit production of high rates of USVs (males/females) or high rates of courtship squeaks (females only) and then was treated with 4-hydroxytamoxifen to drive tdTomato expression in neurons that upregulated Fos in association with that vocalization type, including RAm neurons. Ten days later, subjects were either placed in the same context a second time or were placed in the alternate context (females only). Two hours after this second session, brains were collected and RAm sections were immunostained for Fos expression.

**Figure 3.**
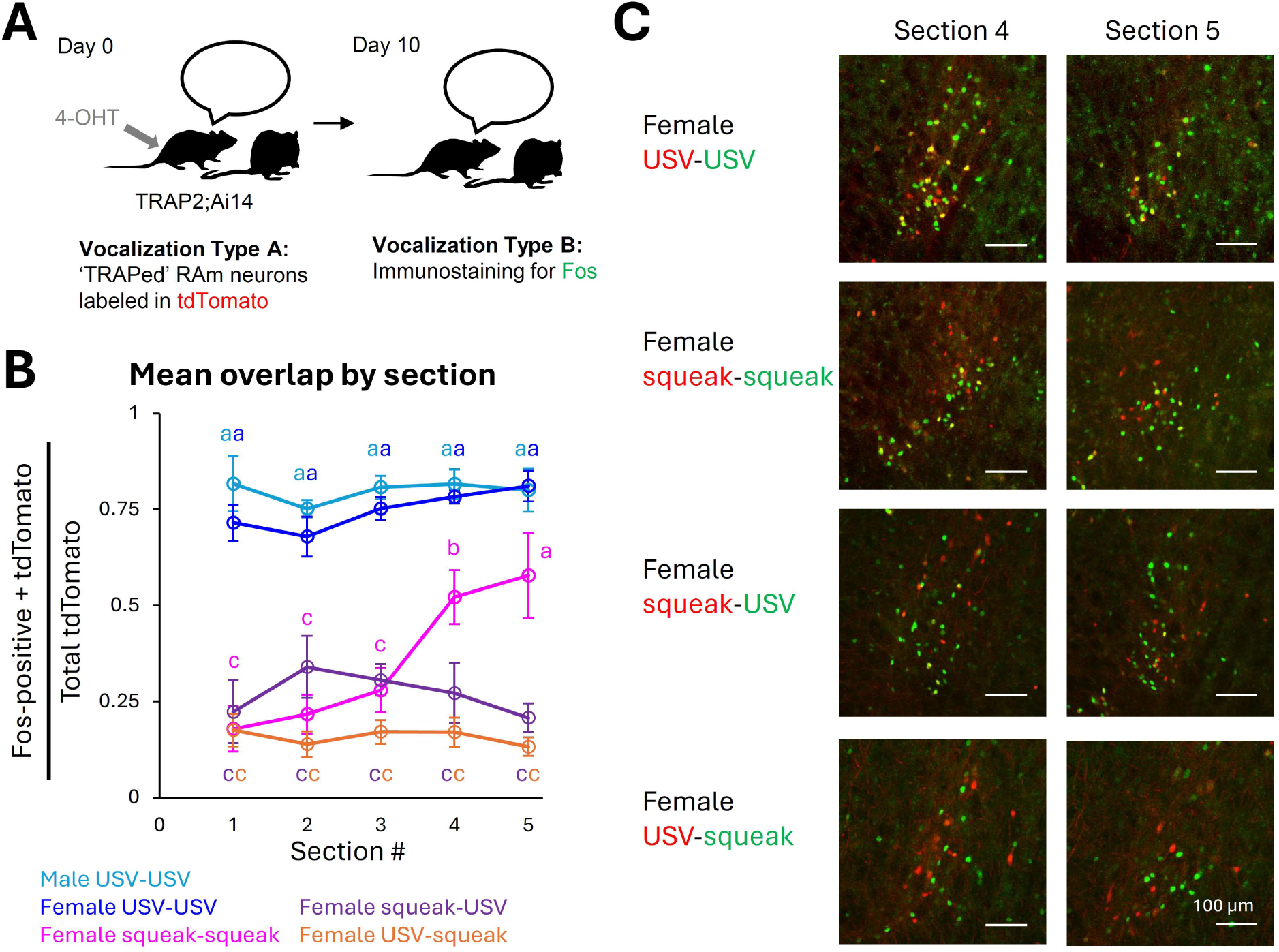
Only a subset of RAm neurons express Fos following the production of both USVs and squeaks. (A) Experimental design. (B) Mean overlap (proportion of tdTomato RAm neurons that are also Fos-positive) is plotted for each group and for each RAm section. Different lowercase letters indicate statistically significant differences between groups for a given RAm section. (C) Representative confocal images showing RAm neurons labeled with tdTomato following behavior 1 (USVs or courtship squeaks) and RAm neurons that are Fos-positive following behavior 2 (USVs or courtship squeaks). Red, tdTomato; green, Fos.

We first evaluated control datasets (n = 5 male USV-USV, n = 6 female USV-USV, and n = 6 female squeak-squeak mice) to ensure that the TRAP2 strategy permits reliable labeling of both USV-related and squeak-related RAm neurons. In the male and female USV-USV groups, a large proportion of tdTomato-expressing RAm neurons that were labeled following the first round of USV production were Fos-positive after the second round of USV production, and mean overlap was high across both rostral and caudal RAm sections (Fig. 3B-C; mean overlap across sections 1-5 for the male USV-USV group is 79.9 ± 8.6%; mean overlap across sections 1-5 for the female USV-USV group is 74.8 ± 6.1%). Heatmap representations of the double-labeled RAm neurons in the female and male USV-USV groups also resemble the pattern of RAm Fos expression following USV production (compare Fig. S3 to Fig. 1B-C), indicating that TRAP2 can efficiently label USV-related RAm neurons. Given that Fos expression associated with squeak production is highest in caudal RAm (Fig. 1D-E) and is only significantly correlated with total squeak time in RAm sections 4-5 (Fig. 1F), we anticipated that the overlap between tdTomato and Fos in the female squeak-squeak group would be highest in caudal RAm. In line with this prediction, we found that mean overlap in the female squeak-squeak group was highest in RAm sections 4-5 (Fig. 3B-C), and heatmap representations of the double-labeled RAm neurons in the squeak-squeak group resemble the pattern of RAm Fos expression following courtship squeak production (compare Fig. S3 to Fig. 1D). We note that the mean overlap in sections 4-5 is lower in the female squeak-squeak group than in the USV-USV groups, possibly because squeak-associated RAm Fos expression is weaker than USV-associated Fos expression (mean overlap across sections 4-5: female squeak-squeak = 54.6 ± 19.6%; male USV-USV = 81.1 ± 9.5%; female USV-USV = 79.9 ± 6.2%). Nonetheless, these data indicate that TRAP2 can reliably label both USV-related and squeak-related ensembles of RAm neurons.

We next tested whether the same RAm neurons are Fos-positive in association with the production of both vocalization types by evaluating overlap in the female squeak-USV group (n = 5) and the female USV-squeak group (n = 6). Notably, overlap in these experimental groups was significantly lower across all RAm sections than the USV-USV control groups and was also lower across caudal RAm sections than the female squeak-squeak control group (Fig. 3C; F(10.76, 61.85) = 4.91, p < 0.001 for interaction effect; one-way ANOVA plus post-hoc Tukey’s HSD for within-section, across-group comparisons only; for each RAm section, significant differences between groups are indicated by different lower-case letters). Together with our Fos datasets, these findings do not support the idea that squeak-related RAm neurons comprise a subset of RAm neurons recruited during USV production. Rather, they suggest a model in which there are at least three types of vocalization-related RAm neurons: (1) RAm-USV neurons, which are recruited during USV production and are distributed throughout rostral and caudal RAm; (2) RAm-squeak neurons, which are recruited during squeak production and are localized in caudal RAm; and (3) RAm-shared neurons, which are recruited during both vocalization types and may be distributed throughout rostral and caudal RAm (Fig. 4).

**Figure 4.**
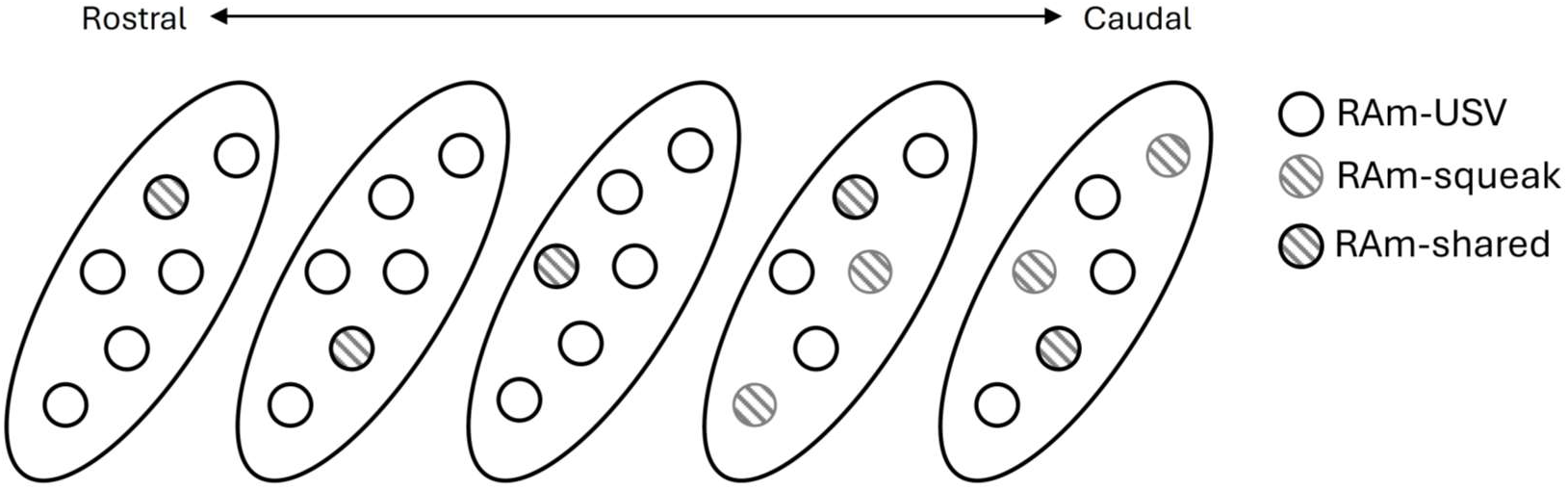
Working model of RAm organization with respect to vocalization type. Our data indicate that RAm may contain USV-related neurons, squeak-related neurons, and shared neurons that are recruited during the production of both vocalization types.

## Discussion

In the current study, we compared patterns of Fos expression in RAm associated with production of USVs by female and male mice and production of courtship squeaks by females. Our across-subjects Fos datasets provide evidence that USV production drives robust Fos expression in both rostral and caudal RAm, whereas female courtship squeak production drives overall weaker Fos expression that is biased toward caudal RAm. Control comparisons indicate that these differences in RAm Fos expression are driven by the production of different vocalization types, rather than by a difference in time spent vocalizing between the USV and courtship squeak groups. Finally, our within-subjects overlap dataset reveals that only a subset of RAm neurons are Fos-positive following the production of both vocalization types, which suggests that the majority of vocalization-related RAm neurons are not recruited in a shared manner across both vocalization types but rather are recruited selectively during either the production of USVs or squeaks.

Our finding that squeak production is associated with fewer Fos-positive RAm neurons than USV production intersects with recent work characterizing the role of *Nts*-expressing RAm neurons in mouse vocal production (Veerakumar et al., 2023). Specifically, Veerakumar et al. found that low frequency optogenetic activation of RAm *Nts* neurons elicits squeak-like (< 20 kHz) vocalizations, while higher frequency optogenetic activation (which presumably drives stronger activation of individual RAm neurons, as well as the activation of greater numbers of RAm neurons) drives the production of vocalizations in the ultrasonic range. These findings are at face value concordant with our observation that USV production is associated with more robust and more spatially extensive RAm Fos expression that courtship squeak production.

Nonetheless, it is somewhat counterintuitive that natural squeak production recruits fewer Fos-positive RAm neurons than the production of USVs. On the one hand, RAm has been implicated in regulating expiratory force and, in turn, vocalization amplitude (i.e., loudness) (Jürgens, 2002; Concha-Miranda et al., 2022; Veerakumar et al., 2023), and so one idea is that increasing RAm activation drives increasing expiratory force in a monotonic fashion and therefore leads to a switch from voiced (i.e., human-audible) to ultrasonic vocalizations. However, past work in rats reported that subglottal pressure is lower during the production of USVs than during the production of squeaks (Riede, 2018). Together with our data showing that only a subset of RAm neurons are Fos-positive following the production of both vocalization types, these findings are difficult to reconcile with the idea that RAm vocalization-related neurons comprise a homogenous population, with increasing activation driving increased expiratory force and therefore a switch from audible to ultrasonic vocalization.

Instead, we hypothesize that Fos-positive RAm neurons associated with the production of USVs vs. squeaks may differ substantially in their connectivity with vocal-respiratory hindbrain regions. Given that the production of rodent USVs relies predominantly on laryngeal but not articulatory mechanisms (Mahrt et al., 2016; Riede et al., 2017; Håkansson et al., 2022), one possibility is that the axonal projections of USV-related RAm neurons are biased toward nucleus ambiguus and spinal expiratory centers, while squeak-related RAm neurons may have particularly strong projections to expiratory centers, in addition to nucleus ambiguus and brainstem articulatory motor neurons. In mammals, RAm has been shown to innervate the spinal cord expiratory centers, nucleus ambiguus, respiration-related hindbrain regions, as well as the trigeminal, facial, and hypoglossal nuclei (Holstege, 1989; Veerakumar et al., 2023). In the zebra finch, RAm neurons that send axonal projections to vocal motor neurons in the hypoglossal nucleus (which in turn innervate the bird’s vocal organ, the syrinx) are spatially intermingled with but distinct from RAm neurons that project to expiratory-related neurons in the thoracic spinal cord (Wild et al., 2009). However, the extent to which projection-defined RAm populations in the mouse are distinct, overlapping, and/or organized topographically remains poorly described.

Our findings highlight the need to characterize heterogeneity within vocalization-related RAm neurons, and to further characterize our Fos-identified populations in terms of their inputs, axonal projections, and expression of molecular markers. Such advances would in turn facilitate the study of these vocalization-related RAm neurons in both females and males.

We also note that there is past work in rats suggesting that RAm neurons differentially regulate the production of different vocalizations. Hartmann and Brecht found that microstimulation of RAm elicited the production of low-frequency (< 30 kHz) USVs but not high frequency (> 30 kHz) USVs, and local cooling of RAm blocked the production of low-frequency USVs elicited by PAG stimulation but did not block PAG-elicited high frequency USVs (Hartmann and Brecht, 2020). Although that study did not evaluate the role of RAm in regulating the production of non-ultrasonic vocalizations, their findings provide additional evidence that RAm neurons participate differentially in the production of different vocalization types.

Past work also explored whether distinct or overlapping populations of PAG neurons regulate the production of different vocalization types. Older work found that partial PAG lesions abolish the production of some vocalization types while leaving others intact, suggesting that different populations of PAG neurons may control the production of different vocalization types (Kelly et al., 1946, 1946; Skultety, 1962; Newman and MacLean, 1982; Jürgens, 1994). In recent work from our group, we used TRAP2 activity-dependent labeling to express caspase in PAG neurons whose activity is critical for the production of USVs. This manipulation blocks the production of USVs in both male and female mice but spares the production of squeaks (including footshock-elicited squeaks in males and females, as well as female courtship squeaks), suggesting that distinct PAG neurons regulate the production of USVs vs. squeaks (Ziobro et al., 2024). The findings of the current study refine our understanding of how regions downstream of the PAG are organized to regulate the production of different vocalization types and suggest that even in the hindbrain, vocalization-related premotor neurons recruited during the production of USVs and squeaks may be largely non-overlapping.

An important limitation of the current study is that the absence of Fos expression does not correspond to the absence of neural activity. Although our results support the conclusion that a larger and more spatially distributed population of RAm neurons is strongly recruited during the production of USVs than during the production of squeaks, our results do not rule out the possibility that some RAm neurons exhibit activity during the production of both vocalization types that fails to drive Fos expression. In vivo measurements and functional manipulations of neural activity in these Fos-defined populations will be important to further characterize these neurons and their contributions to vocal production. Our findings form a foundation to understand how hindbrain circuits are organized to regulate the production of different vocalization types, including in future work that could explore how vocalization-related neurons in RAm and other hindbrain regions are organized to regulate vocal production in different species with diverse vocal repertoires.

## Acknowledgements

We thank the CARE staff for their excellent mouse husbandry, and thanks to Renee Henderson for discussions of the project and manuscript.

