## Supplemental figures for "Production of mouse ultrasonic vocalizations and distress calls is associated with different patterns of Fos expression in the nucleus retroambiguus"

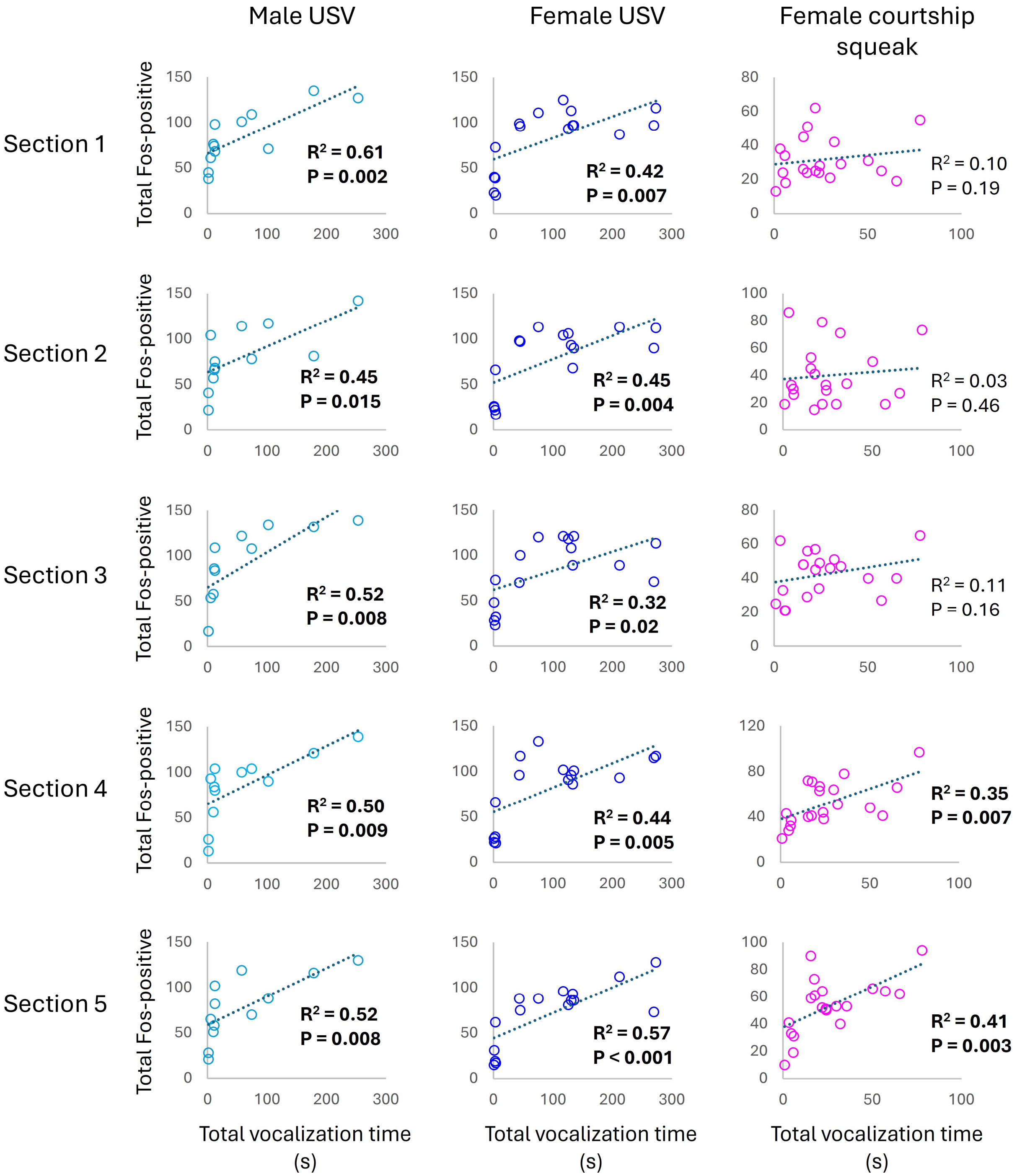


**Figure S1. Relationship between total vocalization time and total Fos-positive RAm neurons.** These plots show the relationship between total vocalization time (in seconds; either USVs or courtship squeaks) vs. total Fos-positive RAm positive neurons, by RAm section (1-5) for each experimental group. Light blue, male USV (n = 12); dark blue, female USV (n = 16);

magenta, female courtship squeak group (n = 20). R-squared values and p values for statistically significant relationships are shown in bold font.

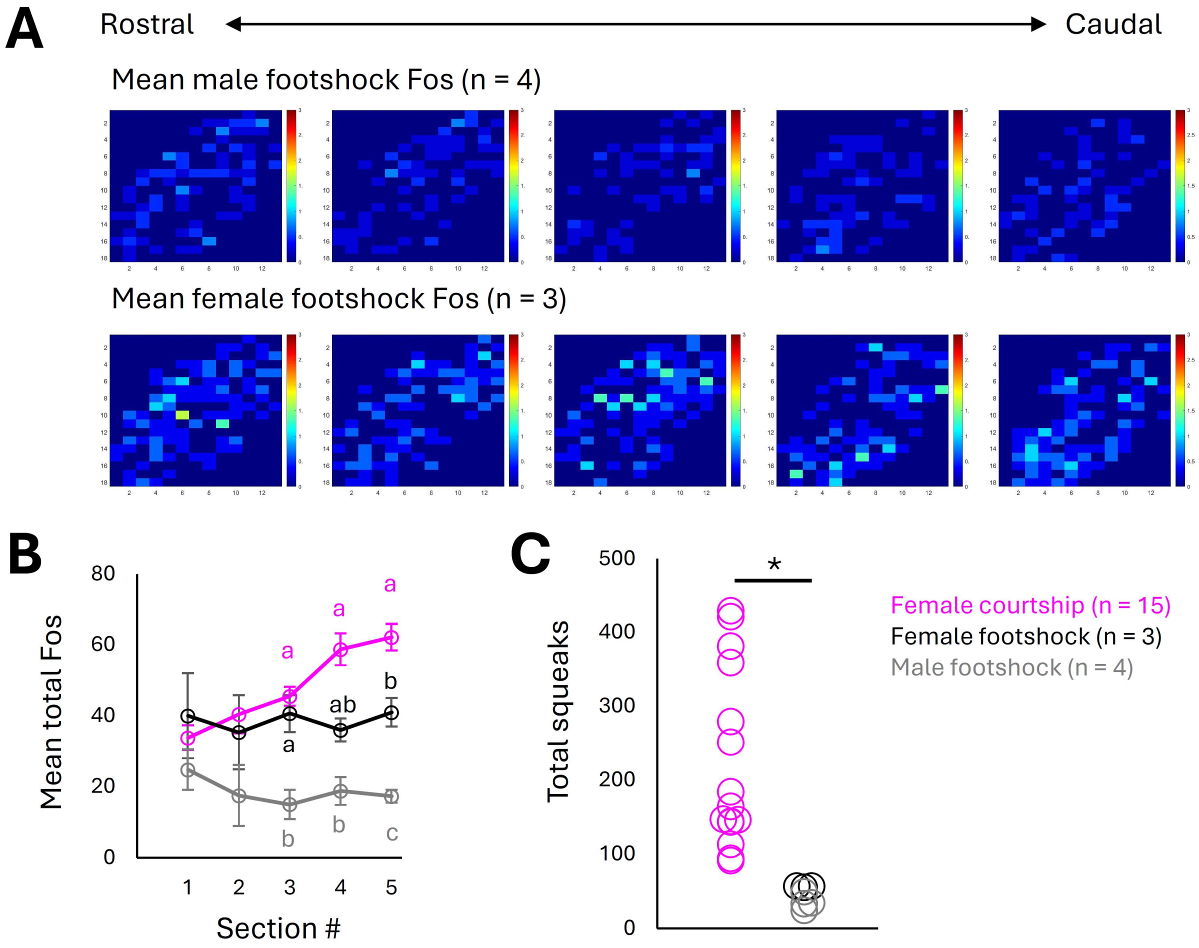


**Figure S2. Comparison of squeak production and RAm Fos expression in the female courtship and male/female footshock groups.** (A) Heatmap representations show mean RAm Fos expression in males and females exposed to a mild footshock paradigm. (B) Mean counts of Fos-positive RAm neurons are shown by section (1-5) in females that produced courtship squeaks (magenta, n = 15), females that received footshocks (black, n = 3), and males that received footshocks (gray, n = 4). Different lowercase letters indicate statistically significant differences between groups for a given RAm section ($F(4,46) = 3.32$, $p = 0.016$ for interaction between group and section; two-way ANOVA with repeated measures on one factor plus Tukey's posthoc for within-section, across-group comparisons only). (C) Females produce significantly greater counts of squeaks during 30-minute courtship interactions with males than do males/females exposed to a mild footshock paradigm ($t(20) = 3.85$, $p < 0.001$; unpaired t-test).

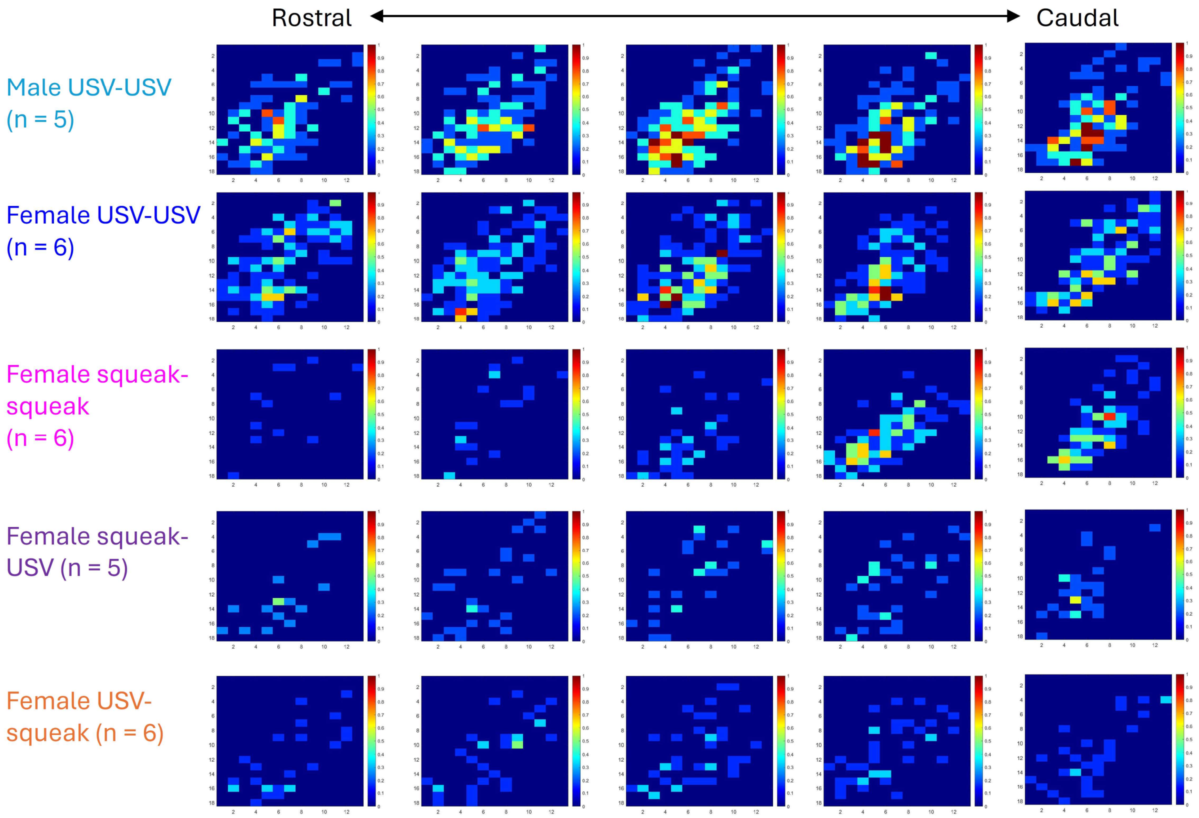


**Figure S3. Heatmap representations of double-labeled RAm neurons (tdTomato-positive and Fos-positive) from RAm overlap experiments.** Heatmaps show the mean distribution of double-labeled RAm neurons. Each row shows a different experimental group, and columns show RAm sections 1-5.
